# Systematic characterization of protein structural features of alternative splicing isoforms using AlphaFold 2

**DOI:** 10.1101/2024.01.30.578053

**Authors:** Yuntao Yang, Yuhan Xie, Zhao Li, Chiamaka S. Diala, Meer A. Ali, Rongbin Li, Yi Xu, Albon Wu, Pora Kim, Sayed-Rzgar Hosseini, Erfei Bi, Hongyu Zhao, W. Jim Zheng

## Abstract

Alternative splicing is an important cellular process in eukaryotes, altering pre-mRNA to yield multiple protein isoforms from a single gene. However, our understanding of the impact of alternative splicing events on protein structures is currently constrained by a lack of sufficient protein structural data. To address this limitation, we employed AlphaFold 2, a cutting-edge protein structure prediction tool, to conduct a comprehensive analysis of alternative splicing for approximately 3,000 human genes, providing valuable insights into its impact on the protein structural. Our investigation employed state of the art high-performance computing infrastructure to systematically characterize structural features in alternatively spliced regions and identified changes in protein structure following alternative splicing events. Notably, we found that alternative splicing tends to alter the structure of residues primarily located in coils and beta-sheets. Our research highlighted a significant enrichment of loops and highly exposed residues within human alternatively spliced regions. Specifically, our examination of the Septin-9 protein revealed potential associations between loops and alternative splicing, providing insights into its evolutionary role. Furthermore, our analysis uncovered two missense mutations in the Tau protein that could influence alternative splicing, potentially contributing to the pathogenesis of Alzheimer’s disease. In summary, our work, through a thorough statistical analysis of extensive protein structural data, sheds new light on the intricate relationship between alternative splicing, evolution, and human disease.

## 1. Introduction

Alternative splicing (AS) is a cellular process that generates multiple mRNAs and protein isoforms for a single gene [1]. This occurs in up to 95% of human genes containing multiple exons [2] and results in a variety of splicing isoforms, including canonical and alternative isoforms. Defined by UniProtKB, canonical isoforms are the most prevalent, most conserved, most expressed and longest isoforms [3]. Splicing isoforms can serve different functions due to varied protein structures and play a crucial role in several biological processes [4]. Alternative splicing is also linked to tumor development [5, 6], which can open new avenues for novel cancer therapies. Computational studies in the field have thus far focused mainly on identifying new transcripts through high-throughput sequencing [7], exploring the role of RNA structure in alternative splicing [8], and investigating alternative splicing signatures of human diseases [9].

In spite of its fundamental importance, very few computational studies have analyzed the changes caused by alternative splicing events at the protein structure level. One study investigated the association between alternative splicing and protein structure evolution and discovered non-trivial alternative isoforms with different functions compared to their canonical isoforms [10]. Another study discovered that the boundaries of alternative splicing sites were enriched with coil and exposed residues [11]. Intrinsically disordered regions were also found in 81% of alternatively spliced regions [12]. However, there are two significant limitations associated with data quality in these studies. First, the protein structural data was obtained from the Protein Data Bank (PDB). While PDB has 167,689 three-dimensional protein structures generated by experimental approaches [13], it covers only 35% of the human proteome. In addition, most of these protein structures are incomplete [3], mainly because experimental approaches cannot obtain structural information of disordered regions. Second, these studies obtained structures of alternative isoforms by sequence alignment to existing structures in PDB, which cannot provide accurate structural data for full-length proteins. These two limitations have precluded systematic investigations into the protein structural changes caused by alternative splicing events.

AlphaFold 2 (AF2) [14], developed by Google DeepMind, significantly outperformed other models in the 14th Critical Assessment of Structural Prediction (CASP14) competition for accurate protein structure prediction. Additionally, DeepMind also constructed the AlphaFold Protein Structure Database (AF2 DB), which includes over 200 million protein structures for about one million species in its latest release [15]. This is a remarkable achievement that has provided us with an unprecedented opportunity to gain structural insights into the alternatively spliced regions. APPRIS, a database for functional annotations of protein isoforms, has incorporated protein structural data from AF2 DB [16], but it has not enabled the detection of structural features of protein isoforms. In this study, we performed a large-scale data analysis on human alternative splicing events using available structural data from PDB, AF2 DB, and our AF2 predictions. We used full-length sequences to generate complete protein structures of canonical and alternative isoforms to address the data quality issues in previous studies. Our comprehensive statistical analysis has identified the protein structural features in alternatively spliced regions and has characterized the structural changes caused by alternative splicing events. This study has been accepted and presented at the International Conference on Intelligent Biology and Medicine (ICIBM 2023) [17].

## 2. Materials and Methods

Figure 1 shows the workflow of our study design, which includes four steps that we have described below.

**Figure 1.**
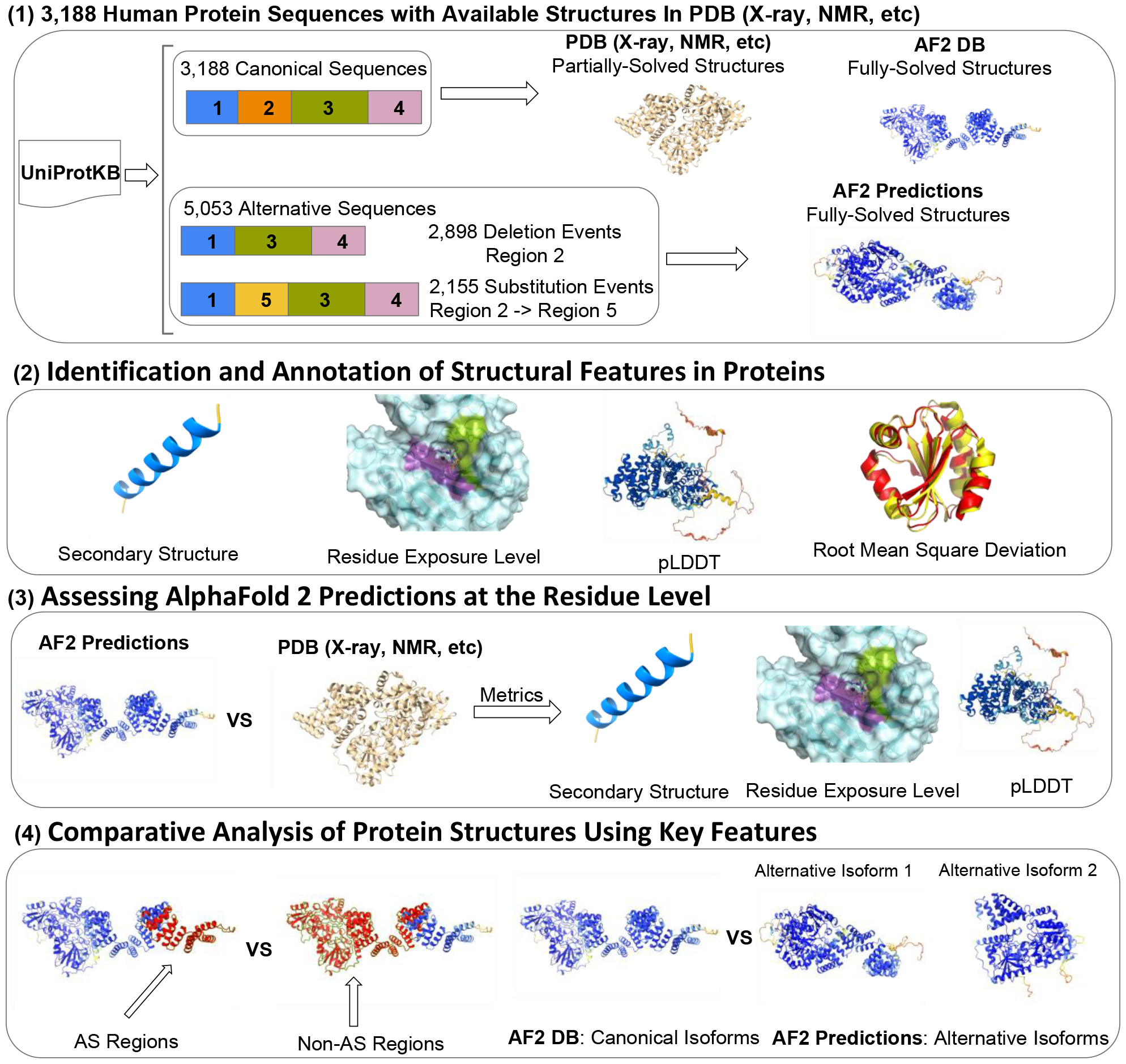
A workflow for the study design.

### 2.1. Step 1: 3,188 Human Protein Sequences with Available Structures In PDB

Raw datasets including the Reviewed Swiss-Prot XML and FASTA sequence datasets, were obtained from UniProtKB [3]. An in-house Python script was used to parse human alternative splicing events and their corresponding isoforms from the Swiss-Prot XML and FASTA datasets. We selected alternative splicing events based on two criteria: (1) alternative splicing events with canonical isoforms, which have available experimental-solved structural data from PDB and (2) only single alternative splicing events in canonical isoforms were selected to minimize the impact of multiple events on one protein structure.

Protein structural data of canonical isoforms were obtained from both PDB [13] and the AF2 DB [18]. PDBrenum was used to renumber residues in PDB structures based on their corresponding UniProtKB sequences [19]. When multiple structures were available for a residue of a protein in PDB, the structure with the highest resolution was selected. Full-length structural data of alternative isoforms is available neither in PDB [13] nor in the AF2 DB [18]. Hence, AF2 was performed to predict their 3D structures [14], which enabled us to construct an in-house AF2 DB. However, AF2’s computing performance is limited in parallel. To improve efficiency, we deployed ParaFold [20], a parallel version of AF2, in a high-performance computing environment. Specifically, the CPU component was deployed on computer clusters at the Texas Advanced Computing Center (TACC), and the GPU component was deployed on Nvidia GPU servers [21] (Figure S1). The max template date for AF2 was set to 2021-07-15.

### 2.2. Step 2: Identification and Annotation of Structural Features in Proteins

We annotated each residue in the isoforms with four types of protein structural features, as outlined in Table S1. Secondary structures and relative accessible surface area (ASA) were obtained using the DSSP program in Biopython [22]. The secondary structures were categorized into both 8-class and 3-class categories, and residue exposure levels were established based on the relative ASA range. Additionally, the residue-level confidence scores (pLDDTs) of predicted structures were obtained from ‘result_model_*.pkl’ in output files of AF2, to evaluate AF2 performance in different protein structural regions. Lastly, for quantifying the structural similarity before and after alternative splicing events, the Root-mean-square deviation (RMSD) was calculated on 3D coordinates of alpha carbons [23]. It is important to note that this metric was not included in previous studies due to the lack of full-length protein structural data.

### 2.3. Step 3: Assessing AlphaFold 2 Predictions at the Residue Level

We acquired data on 8-class and 3-class secondary structures, along with residue exposure levels, for both canonical isoforms (sourced from PDB and AF2 DB) and alternative isoforms (derived from our in-house AF2 DB). To assess the accuracy of AF2’s predictions for canonical isoforms, we calculated the consistency rates for secondary structures and residue exposure levels by comparing overlapping residues between the AF2 predictions and the PDB. For evaluating the predictions of alternative isoforms at in-house AF2 DB, we extracted pLDDTs and compared them with the predictions for canonical isoforms from the AF2 DB.

While AF2 provides precise structure predictions, there are instances where its confidence in predicting certain protein regions is limited. To address this limitation, we took two measures in this study to mitigate this limitation. We evaluated AF2’s secondary structure predictions in very low-confidence regions, defined as those with pLDDT scores below 50. This evaluation involved a comparative analysis with Jpred4 [24], a well-established neural network-based predictor of 3-class secondary structures. For the Jpred predictions, we specifically selected protein sequences of canonical isoforms that meet the following criteria: their length is less than 800 amino acids (AA), and over 50% of their residues fall into the category of very low confidence according to AF2 predictions. This selection was made, taking into account the limitation of Jpred4 in handling protein sequences of up to 800 AA.

### 2.4. Step 4.a: Alternative Splicing Regions vs. Non-Alternative Splicing Regions

We classified the residues in each isoform into three categories based on their position: *i)* substitution regions, *ii)* deletion regions, and *iii)* non-alternative splicing regions. For canonical isoforms with multiple corresponding alternative isoforms or alternative splicing events, residues not belonging to any alternative splicing regions were classified as non-alternative splicing regions. To determine the enrichment of secondary structures and residue exposure levels, we performed Chi-squared tests in substitution regions and deletion regions, which resulted in three sets of comparison: (1) substitution regions vs. non-alternative splicing regions in canonical isoforms (Comparison 1 in Figure 2A), (2) substitution regions vs. non-alternative splicing regions in alternative isoforms (Comparison 2 in Figure 2A) and (3) deletion regions vs. non-alternative splicing regions in canonical isoforms (Comparison 3 in Figure 2A). The p-values were adjusted with the Benjamini-Hochberg procedure to control the false discovery rate (FDR) in Chi-squared tests.

**Figure 2.**
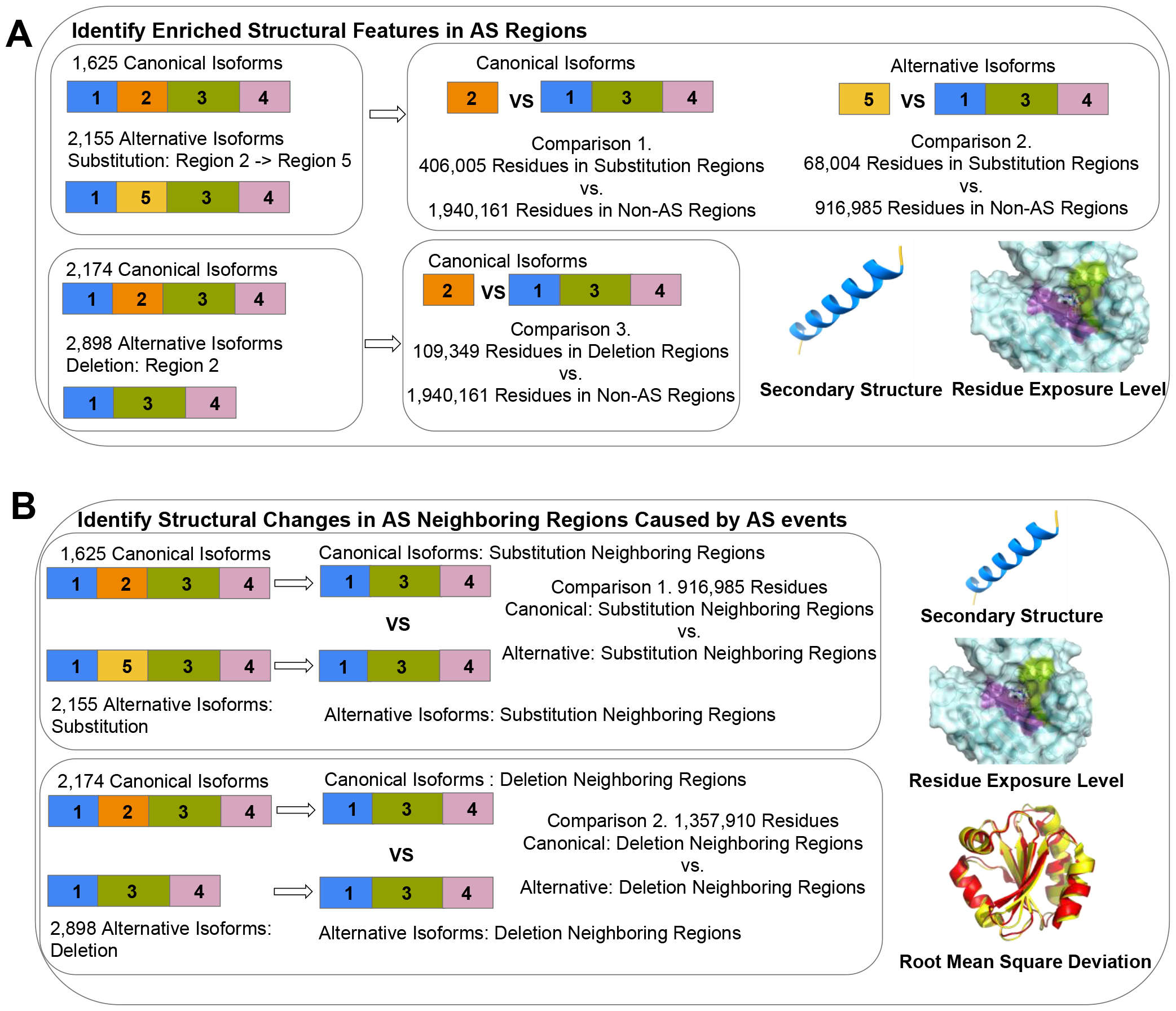
Comparative Analysis of Protein Structures Focusing on Structural Features. (A) Three comparisons for identifying enriched structural features in alternative splicing regions. (B) Two comparisons for identifying structural changes in alternative splicing neighboring regions caused by alternative splicing events.

### 2.5. Step 4.b: Canonical Isoforms vs. Alternative Isoforms

To evaluate the structural changes caused by alternative splicing events, we compared the canonical isoforms with their corresponding alternative isoforms for each residue in the regions surrounding alternative splicing events. We examined changes in secondary structures and residue exposure levels by using McNemar tests, which enabled us to compare the canonical isoforms with alternative isoforms for each residue in substitution neighboring regions (Comparison 1 in Figure 2B) and deletion neighboring regions (Comparison 2 in Figure 2B). The minimum p-value was set at 1e-99 for visualization purposes, and p-values were adjusted with the Benjamini-Hochberg procedure to control the false discovery rate (FDR) in the McNemar tests. To further assess the structural changes in the regions surrounding alternative splicing events, we also used RMSD. We performed the unpaired Wilcoxon signed-rank test to determine the difference between substitution events and deletion events regarding structural changes.

## 3. Results

### 3.1. Systematic characterization of protein structural features of alternatively spliced regions

#### 3.1.1.The Alternative Splicing Events and Isoforms

We collected a comprehensive catalogue of alternative splicing events and isoforms from the UniProtKB database, which contains 28,063 human alternative splicing events. However, for only 14,645 events with canonical isoforms, structural data in PDB are available. After selecting single alternative splicing events in canonical isoforms, 5,716 events were included in our study. Finally, we ended up with 3,188 structures for the canonical isoforms and 5,053 structures for alternative isoforms, which are available in our in-house AF2 DB. Hence, our study finally includes 2,155 substitution events and 2,898 deletion events. Figure S2 displays the length distributions of deletion and substitution regions in canonical isoforms, with most lengths being under 100 amino acids (AA). The concise nature of these regions in the figure indicates that our conclusions are primarily based on individual amino acid properties, minimizing the influence of fragment length.

#### 3.1.2. Assessing AlphaFold 2 Predictions at the Residue Level

To evaluate AF2’s effectiveness in predicting canonical isoforms, we analyzed the consistency rates of secondary structures and residue exposure levels by comparing overlapping between the PDB and AF2 DB. The results showed that the AF2 DB attained an 87.36% consistency rate for 8-class secondary structures and 92.53% for 3-class secondary structures. However, the consistency rate for residue exposure levels was notably lower at 66.59%. This disparity suggests that while AF2 excels in predicting secondary structures, likely due to its utilization of PDB structures as templates, its accuracy in predicting residue exposure levels is limited, primarily because it captures only partial domain-packing information.

Further analysis of alternative isoforms indicated that AF2’s predictions for these isoforms are comparably accurate to those for canonical isoforms within the AF2 DB, as illustrated in Figure 3A. Additionally, we conducted a comparative study between AF2 and Jpred4 [24], specifically focusing on 3-class secondary structure predictions. From a pool of 3,534 canonical isoforms, 168 were chosen based on criteria specified in section 2.3 of the Methods. Our findings showed that the overall consistency rate for all residues between AF2 and Jpred4 is 85%. Interestingly, for residues with very low confidence, the consistency rate between AF2 and Jpred4 increases to 95%. Consequently, the study demonstrates that both AF2-predicted canonical and alternative isoforms maintain high levels of accuracy.

**Figure 3.**
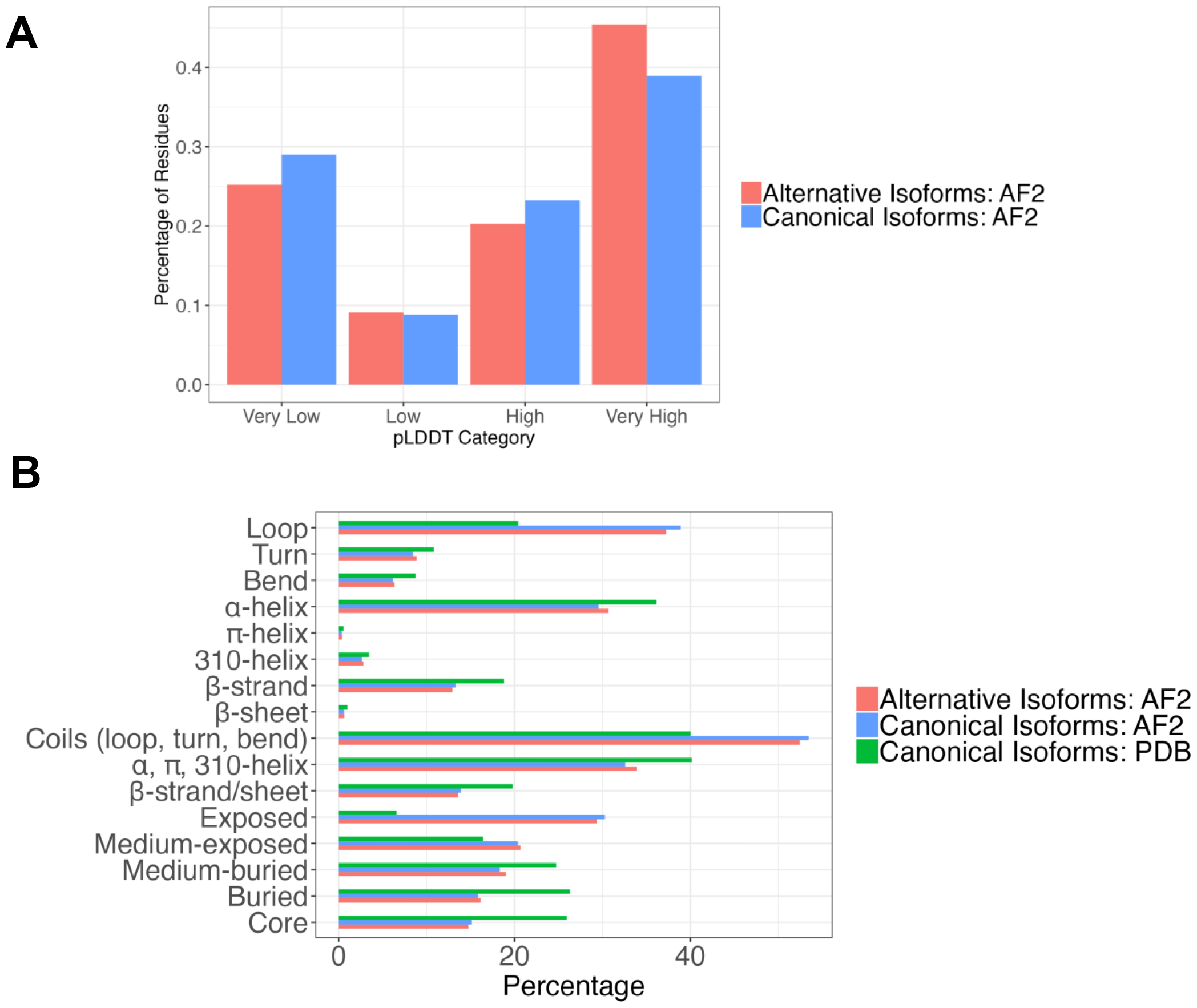
Assessment of AF2 predictions. (A) Compare the pLDDT of alternative isoforms (in-house AF2 DB) with canonical isoforms (AF2 DB). (B) Percentages of protein structural features for canonical isoforms and alternative isoforms.

However, PDB has a lower percentage of loop/coil secondary structures than the AF2 DB and the in-house AF2 DB (Figure 3B), as residues in loops or coils, could be disordered regions that lack well-defined structures in PDB. Additionally, the percentage of exposed residues was found to be lower in PDB, while core and buried residues had higher percentages (Figure 3B). It is challenging for the experimental methods in PDB data to provide structural information for residues in loops or coils and exposed regions.

In summary, for the subsequent analyses, we decided to use the AF2 DB as the source for canonical isoform structures, because AF2 predictions provide unbiased structural data that includes more isoforms and cover protein sequences more thoroughly as compared to PDB, which provides only partial coverage. It is important to note that we only utilized AF2 to supplement the gaps in PDB structures. This decision was based on our selection of canonical isoforms that possess structures in both AF2 and PDB databases. For alternative isoforms, we performed AF2 predictions in-house. Those alternative isoforms with a single alternative splicing event have sequences akin to canonical isoforms, enabling AF2 to identify available structural templates for higher-confidence predictions. In addition, our comparative analysis with Jpred4 further validated the reliability of AF2 predictions in secondary structures, reinforcing our decision to use AF2 predictions in our study.

#### 3.1.3. Alternative vs. Non-Alternative Splicing Regions

We analyzed the structural features of over 5,000 substitution and deletion events in canonical isoforms and over 2,000 substitution events in alternative isoforms, to identify structural features enriched in alternative splicing regions. Figure 4A shows that substitution and deletion regions in canonical isoforms are enriched with exposed residues and coils/loop secondary structures. Likewise, substitution regions in alternative isoforms also have a high prevalence of these structural features. The presence of loops and coils in alternative splicing regions indicates that these regions are mostly intrinsically disordered. This supports the hypothesis that alternative splicing regions are characterized by intrinsically disordered and exposed features, which contribute to protein diversity without affecting the core structural regions, as previously reported in a study of 46 protein structures [12].

**Figure 4.**
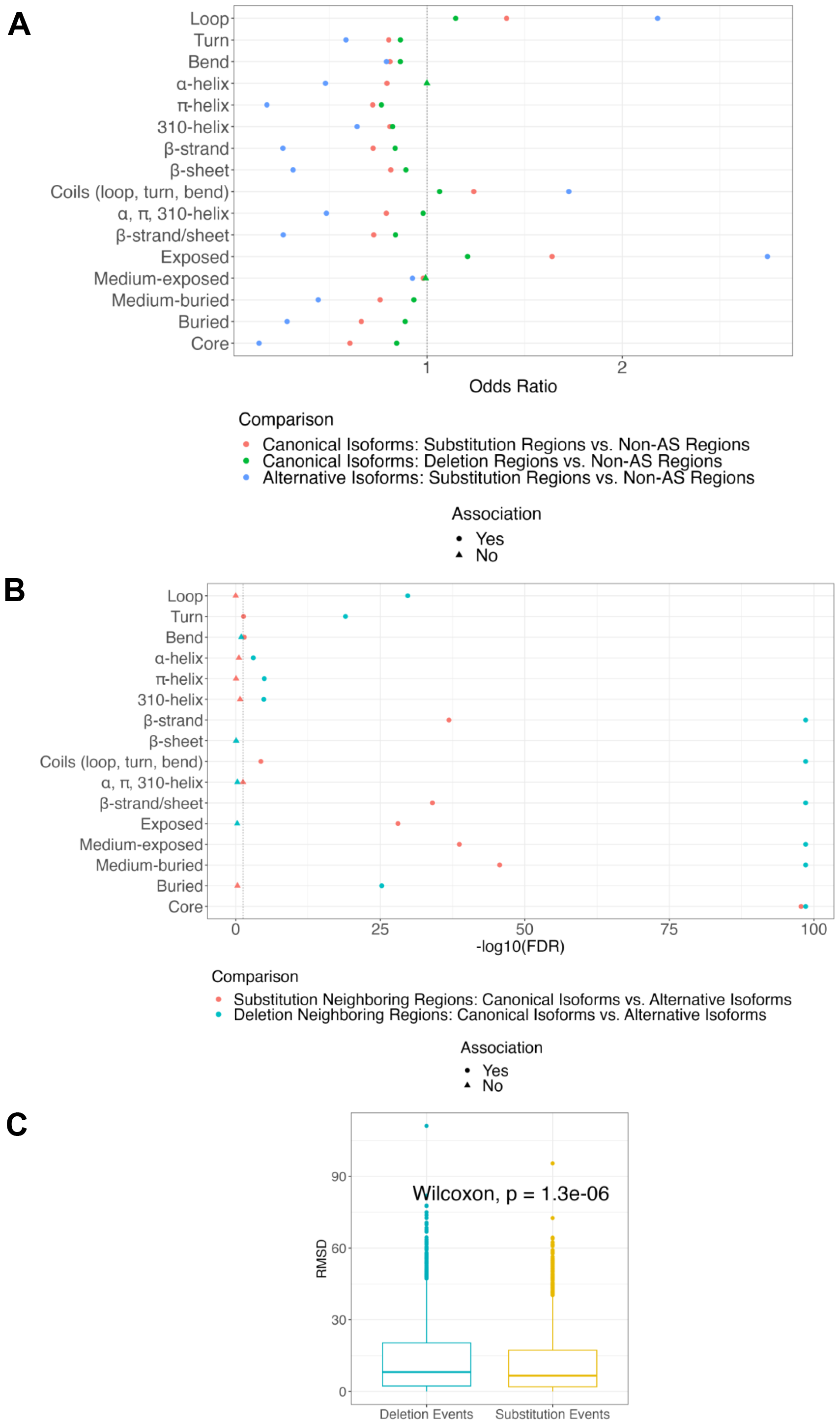
Comparative analysis of protein structures. (A) Enrichment of structural features in alternative splicing regions (significant enrichment: FDR < 0.05 and Odds Ratio > 1). (B) Structural changes in alternative splicing neighboring regions (significant change: FDR < 0.05). (C) Comparison of RMSD between substitution neighboring regions and deletion neighboring regions.

#### 3.1.4. Canonical vs. Alternative Isoforms

Structural changes resulting from alternative splicing events were evaluated by comparing canonical and alternative isoforms in the alternative splicing neighboring regions. We analyzed over 3000 canonical isoforms and over 5000 corresponding alternative isoforms to study changes in structural features after alternative splicing events. In substitution neighboring regions, only 3 out of 8 secondary structure features were significantly altered after substitution events, while buried residues remained unchanged (Figure 4B). For deletion neighboring regions, 6 out of 8 secondary structures showed significant changes after deletion events, while exposed residues remained unchanged (Figure 4B). Compared to substitution events, deletion events resulted in greater changes to secondary structures in neighboring regions and tended to alter core structures in these regions. This suggests that deletion events might have a greater impact on protein function compared to substitution events. Coils and beta strands/sheets were found to be unstable after both substitution and deletion events. Coils are mainly intrinsically disordered regions that include loops, turns, and bends, which are unstable after alternative splicing events because of the lack of fixed structures. The beta-strand is a secondary structure formed by hydrogen bonds, which can be easily disrupted by alternative splicing events [10]. Thus, isoforms that are enriched with coils and beta strands/sheets tend to have novel functions after alternative splicing events due to significant structural changes.

We collected ∼2,800 substitution neighboring regions and ∼4,600 deletion neighboring regions with available structural data for both canonical and alternative isoforms. The RMSDs were calculated by comparing the canonical isoforms with their corresponding alternative isoforms in their alternative splicing neighboring regions. The results showed that deletion events have significantly higher RMSDs than substitution events (Figure 4C), suggesting that deletions cause more significant changes to the structure of these regions compared to substitutions. This is consistent with the results obtained from the analysis of secondary structures (Figure 4B).

### 3.2. Case Study 1: Septin-9 Protein and Its Alternative Splicing Events

Intrinsically disordered regions are frequently characterized as loops, which are significantly enriched in alternative splicing regions. To explore the relationship between alternative splicing and loops, we proposed a hypothesis that intrinsically disordered regions in protein structures drive alternative splicing events during evolution. To test this hypothesis, we selected the Septin-9 protein, a crucial protein involved in cytokinesis and many other biological processes. This protein has various homologs found in species ranging from fungi to mammals (Table S2). It produces multiple alternative isoforms in several species including *Xenopus tropicalis, Mus musculus, Rattus norvegicus, Homo sapiens*, and *Bos taurus*, while no alternative isoforms are found in *Danio rerio, Gallus gallus*, and all fungi. Figure 5A presents the AF2-predicted structure of the Septin-9 protein in *Saccharomyces cerevisiae*, whereas Figure 5B depicts the corresponding AF2-predicted structure of the Septin-9 protein in *Homo sapiens*.

**Figure 5.**
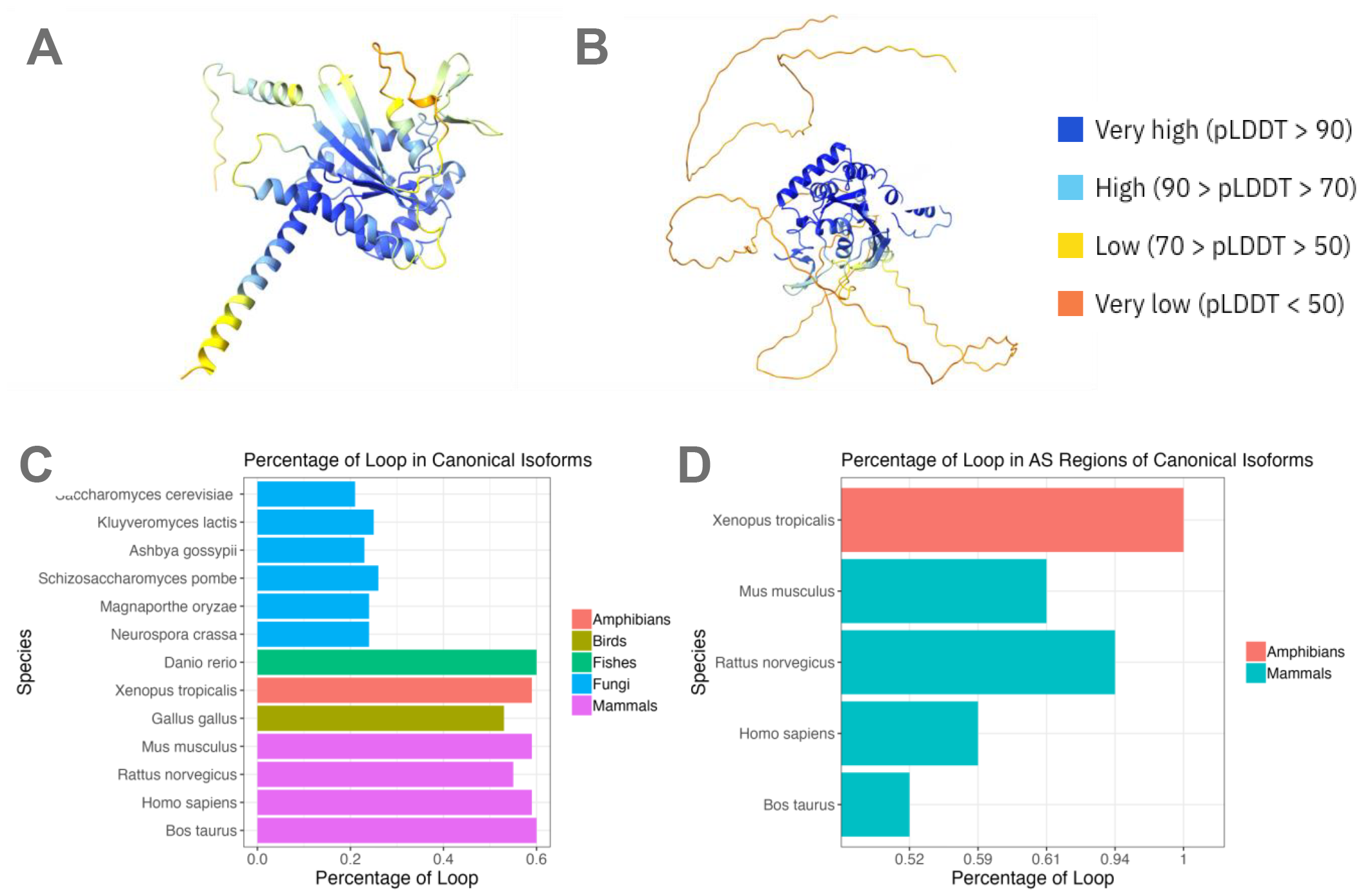
(A) 3D structures for Septin-9 proteins of *Saccharomyces cerevisiae*. (B) 3D structures for Septin-9 proteins of *Homo sapiens*. (C) Bar plot for percentage of the loop in full sequences of canonical isoforms. (D) Bar plot for percentage of the loop in alternative splicing regions of canonical isoforms.

We analyzed secondary structures and alternative splicing regions of Septin-9 homologs in each species. The proportion of loop regions exhibited a significant increase, rising from around 20% in fungi to more than 50% in other complex species, as illustrated in Figure 5C. Conversely, all other secondary structures experienced a decrease, as documented in Table S3. Furthermore, over 50% of residues in alternative splicing regions are found in loop structures in *Xenopus tropicalis* and mammals (Figure 5D). Figures 5A and 5B display the significant structural changes of Septin-9 from *Saccharomyces cerevisiae* to *Homo sapiens*, specifically in loop regions. While the AF2 predictions for loop regions in Septin-9 primarily exhibit a pLDDT of less than 50, a pLDDT value below 50 is a reliable indicator of disordered regions. These results support our hypothesis that intrinsically disordered regions potentially contribute to protein diversity in complex species by promoting alternative splicing events.

### 3.3. Case Study 2: Missense Mutations in Alternative Splicing Regions of Tau Protein

The human Tau protein is an intrinsically disordered protein that binds to microtubules in neurons [25]. The binding dysfunction to microtubules of the Tau protein is a hallmark of Alzheimer’s disease [26]. Alternative splicing events in the Tau protein produce isoforms with varying numbers of N-terminal projection domains (N regions) and microtubule-binding regions (R regions). There are six different isoforms of the human Tau protein found in the central nervous system, including three 4R isoforms (2N4R Tau, 1N4R Tau, 0N4R Tau) and three 3R isoforms (2N3R Tau, 1N3R Tau, and 0N3R Tau) [27]. The canonical 2N4R Tau isoform contains four R regions, including R1, R2, R3, and R4. R2, the alternative splicing region, is absent in the 3R isoforms. The ratio of 4R and 3R isoforms in the normal adult brain is approximately 1:1, but this ratio becomes imbalanced in the neurodegenerative brain [28]. We hypothesize that missense mutations in the R2 region of the 2N4R Tau might alter the protein structure from random coils to more structured regions, leading to changes in alternative splicing events and the imbalance of the 4R and 3R isoform ratio.

Missense mutations in the R regions of the 2N4R Tau isoform were collected from the ALZFORUM database [29]. Using the AF2 program [14], we predicted the protein structures of the wild-type and mutated 2N4R isoforms. Table 1 shows the number of mutations that alter secondary structures. In the R1 region, there are two missense mutations that cause structural changes, including I260V (loop to bend) and G273R (loop to turn). However, these mutations do not stabilize the structure into a fixed state, such as a helix or strand. In the R2 region, there are four missense mutations that cause structural changes, including L284R (loop to 3-10 helix), S285R (turn to 3-10 helix), N296D (loop to turn), and N296H (loop to turn). The mutations L284R and S285R transformed residues 284 to 286 from coils to helix structures. No structural changes were detected in the missense mutations in the R3 and R4 regions. Detailed secondary structural changes can be found in Table S4.

**Table 1.** Numbers of missense mutations in R regions.

| Regions | Number of Missense Mutations<br>(Secondary Structure Change) | Number of Missense Mutations<br>(No Secondary Structure Change) |
| --- | --- | --- |
| R1 | 2 | 4 |
| R2 | 4 | 8 |
| R3 | 0 | 11 |
| R4 | 0 | 10 |

The L284R mutation was identified in a Caucasian family [30], and the S285R mutation was detected in a Japanese man [31], both presenting with progressive supranuclear palsy, a condition characterized by Tau deposition in the brain, similar to Alzheimer’s disease. While L284R has not been confirmed as a pathogenic mutation of Alzheimer’s disease through functional studies, it is still likely to be pathogenic, because several pathogenic mutations have been reported at the same codon [32]. S285R, on the other hand, has been confirmed as a novel pathogenic mutation through an exon-trapping analysis [31]. For the confidence of predictions, the R2 region of the Tau protein, encompassing residues 284-286, undergoes mutation and exhibits a pLDDT value ranging between 50 and 70 for the wildtype, L284R, and S285R variants. Although this range is not exceptionally high, it is still noteworthy for conclusion, particularly when considering that mutations in other R regions did not yield significant structural alterations (Table 1). This evidence supports the hypothesis that missense mutations in the R2 regions are likely to cause structural changes in the protein, which disrupt alternative splicing events and lead to an imbalanced ratio of 4R and 3R isoforms.

## 4. Discussion

The lack of full-length protein structural data in PDB has prohibited a detailed understanding of alternative splicing at the protein structure level. With the advent of AF2, a breakthrough in structural biology, we now have the opportunity to comprehensively decode alternative splicing events. In this study, we utilized AF2 to predict the structures of splicing isoforms and applied statistical methods to provide new insights into the structural features of alternative splicing. Our results, based on a large amount of high-quality protein structural data, not only provide solid statistical evidence but also validate several hypotheses related to alternative splicing events.

Our analyses revealed that alternative splicing in human proteins occurs frequently in intrinsically disordered and exposed regions, which supports the idea that alternative splicing contributes to protein diversity without disrupting core functional structures [12]. The presence of intrinsically disordered regions may also play a role in the evolution of alternative splicing events. While single-cell eukaryotes like *Schizosaccharomyces pombe* lack alternative splicing events [33], up to 95% of human genes exhibit alternative splicing [2]. The Septin-9 protein case study suggests that intrinsically disordered regions in proteins may have facilitated alternative splicing events over the course of evolution. This hypothesis was supported by our observations including the gradual increase of loop residues from single-cell eukaryotes to more complex species, and also the significant enrichment of loop residues in alternative splicing regions of multicellular eukaryotes. Thus, our analysis raises the possibility that intrinsically disordered regions may have played a key role in protein evolution by promoting protein diversity. Additionally, the use of AF2 in predicting protein structures for mutated sequences offers the opportunity to investigate the relationship between alternative splicing events and missense mutations. Our case study of the Tau protein revealed that two missense mutations in the alternative splicing region caused a change from coils to more structured regions, potentially increasing the risk of Alzheimer’s disease by altering the ratio of 4R to 3R isoforms. This corroborates former studies, which have established that missense mutations in the CFTR gene lead to abnormal alternative splicing events and cause cystic fibrosis [34].

Our study can be extended from two directions. Firstly, we have only focused on human genes, and so further research is needed to examine alternative splicing events in other multicellular eukaryotes, such as mice and rats, to gain a more holistic evolutionary insight. Secondly, the predictions made by AF2 are based on existing protein crystal structures that are largely from laboratory experiments, rather than in vivo conditions. Addressing this challenge is important, especially because protein folding can be altered by various factors in vivo, such as post-translational modifications, and hence studying intrinsically disordered proteins using AF2 may not accurately reflect such changes. While our study encountered challenges stemming from the low-confidence predictions of AF2, it’s important to note that AF2 is still at the forefront as a computational tool for bridging the structural information void. To address its low-confidence predictions, we focused on proteins with structures available in both PDB and AF2 we also evaluated AF2’s secondary structure predictions in low-confidence regions by comparing them with Jpred4 [24]. As a result, AF2 primarily complemented missing information in PDB rather than predicting completely novel structures. Furthermore, in these particularly uncertain regions, AF2 demonstrated a consistent alignment with Jpred4’s predictions.

In summary, we have taken a significant step towards comprehensively compiling protein structural features of alternative splicing events in human that can be further extended to other species and in vivo conditions in future studies.

## Acknowledgments

This work is partly supported by the Cancer Prevention and Research Institute of Texas through grant RP170668, the National Institutes of Health (NIH) through grant R01AG066749, 1U24MH130988-01, 5R56AG069880-02, and the Department of Defense (DoD) through grant W81XWH-22-1-0164 to WJZ. The authors acknowledge the Texas Advanced Computing Center (TACC) at The University of Texas at Austin (http://www.tacc.utexas.edu) for providing HPC and database resources.

## References

1. Stamm, S., et al., Function of alternative splicing. Gene, 2005. 344: p. 1–20.

2. Pan, Q., et al., Deep surveying of alternative splicing complexity in the human transcriptome by high-throughput sequencing. Nature genetics, 2008. 40(12): p. 1413–1415.

3. UniProt: the universal protein knowledgebase in 2021. Nucleic acids research, 2021. 49(D1): p. D480–D489.

4. Gallego-Paez, L.M., et al., Alternative splicing: the pledge, the turn, and the prestige. Human genetics, 2017. 136(9): p. 1015–1042.

5. Kahles, A., et al., Comprehensive analysis of alternative splicing across tumors from 8,705 patients. Cancer cell, 2018. 34(2): p. 211–224. e6.

6. Urbanski, L.M., N. Leclair, and O. Anczuków, Alternative-splicing defects in cancer: Splicing regulators and their downstream targets, guiding the way to novel cancer therapeutics. Wiley Interdisciplinary Reviews: RNA, 2018. 9(4): p. e1476.

7. Jiang, W. and L. Chen, Alternative splicing: Human disease and quantitative analysis from high-throughput sequencing. Computational and Structural Biotechnology Journal, 2021. 19: p. 183–195.

8. Xu, B., Y. Meng, and Y. Jin, RNA structures in alternative splicing and back-splicing. Wiley Interdisciplinary Reviews: RNA, 2021. 12(1): p. e1626.

9. Mao, S., et al., Survival-associated alternative splicing signatures in esophageal carcinoma. Carcinogenesis, 2019. 40(1): p. 121–130.

10. Birzele, F., G. Csaba, and R. Zimmer, Alternative splicing and protein structure evolution. Nucleic acids research, 2008. 36(2): p. 550–558.

11. Wang, P., et al., Structural genomics analysis of alternative splicing and application to isoform structure modeling. Proceedings of the National Academy of Sciences, 2005. 102(52): p. 18920–18925.

12. Romero, P.R., et al., Alternative splicing in concert with protein intrinsic disorder enables increased functional diversity in multicellular organisms. Proceedings of the National Academy of Sciences, 2006. 103(22): p. 8390–8395.

13. Berman, H.M., et al., The protein data bank. Nucleic acids research, 2000. 28(1): p. 235–242.

14. Jumper, J., et al., Highly accurate protein structure prediction with AlphaFold. Nature, 2021. 596(7873): p. 583–589.

15. Callaway, E., ‘The entire protein universe’: AI predicts shape of nearly every known protein. Nature, 2022. 608(7921): p. 15–16.

16. Rodriguez, J.M., et al., APPRIS: selecting functionally important isoforms. Nucleic acids research, 2022. 50(D1): p. D54–D59.

17. Yang, Y., et al., ICIBM Program Book, Page 60, https://icibm2023.iaibm.org/Schedule_files/ICIBM2023%20Program_book_final.pdf, in International Conference on Intelligent Biology and Medicine. July 16th - 19th, 2023: Tempa, FL.

18. Varadi, M., et al., AlphaFold Protein Structure Database: massively expanding the structural coverage of protein-sequence space with high-accuracy models. Nucleic acids research, 2022. 50(D1): p. D439–D444.

19. Faezov, B. and R.L. Dunbrack Jr, PDBrenum: A webserver and program providing Protein Data Bank files renumbered according to their UniProt sequences. Plos one, 2021. 16(7): p. e0253411.

20. Zhong, B., et al. Parafold: Paralleling alphafold for large-scale predictions. in International Conference on High Performance Computing in Asia-Pacific Region Workshops. 2022.

21. Yang, Y., et al., AlphaFold 2 Monomer: Deployment in an HPC Environment. TACCSTER 2022 Proceedings, 2022.

22. Kabsch, W. and C. Sander, Dictionary of protein secondary structure: pattern recognition of hydrogen-bonded and geometrical features. Biopolymers: Original Research on Biomolecules, 1983. 22(12): p. 2577–2637.

23. Maiorov, V.N. and G.M. Crippen, Significance of root-mean-square deviation in comparing three-dimensional structures of globular proteins. Journal of molecular biology, 1994. 235(2): p. 625–634.

24. Drozdetskiy, A., et al., JPred4: a protein secondary structure prediction server. Nucleic acids research, 2015. 43(W1): p. W389–W394.

25. Skrabana, R., J. Sevcik, and M. Novak, Intrinsically disordered proteins in the neurodegenerative processes: formation of tau protein paired helical filaments and their analysis. Cellular and molecular neurobiology, 2006. 26(7): p. 1083–1095.

26. Barbier, P., et al., Role of tau as a microtubule-associated protein: structural and functional aspects. Frontiers in aging neuroscience, 2019. 11: p. 204.

27. Ballatore, C., V.M.-Y. Lee, and J.Q. Trojanowski, Tau-mediated neurodegeneration in Alzheimer’s disease and related disorders. Nature reviews neuroscience, 2007. 8(9): p. 663–672.

28. Gasparini, L., B. Terni, and M.G. Spillantini, Frontotemporal dementia with tau pathology. Neurodegenerative diseases, 2007. 4(2-3): p. 236–253.

29. Kinoshita, J. and T. Clark, Alzforum. Neuroinformatics, 2007: p. 365–381.

30. Rohrer, J.D., et al., Novel L284R MAPT mutation in a family with an autosomal dominant progressive supranuclear palsy syndrome. Neurodegenerative Diseases, 2011. 8(3): p. 149–152.

31. Ogaki, K., et al., Analyses of the MAPT, PGRN, and C9orf72 mutations in Japanese patients with FTLD, PSP, and CBS. Parkinsonism & related disorders, 2013. 19(1): p. 15–20.

32. Rossi, G. and F. Tagliavini, Frontotemporal lobar degeneration: old knowledge and new insight into the pathogenetic mechanisms of tau mutations. Frontiers in aging neuroscience, 2015. 7: p. 192.

33. Barrass, J.D. and J.D. Beggs, Splicing goes global. TRENDS in Genetics, 2003. 19(6): p. 295–298.

34. Garcia-Blanco, M.A., A.P. Baraniak, and E.L. Lasda, Alternative splicing in disease and therapy. Nature biotechnology, 2004. 22(5): p. 535–546.

