## Supplemental Figures and Tables for "Systematic characterization of protein structural features of alternative splicing isoforms using AlphaFold 2"

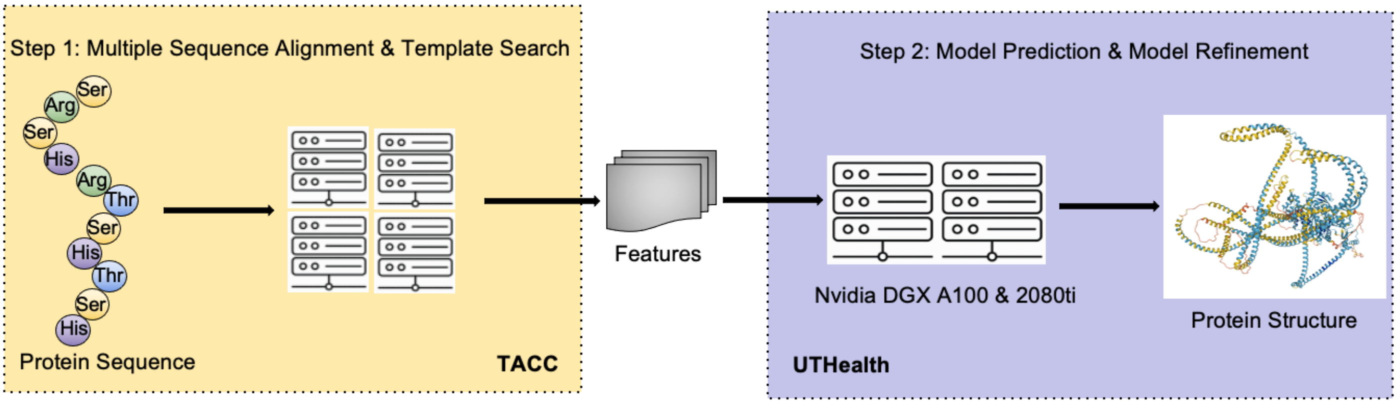


Supplemental Figure S1. Employ high-performance computing to efficiently predict the protein structures of alternative splicing isoforms.

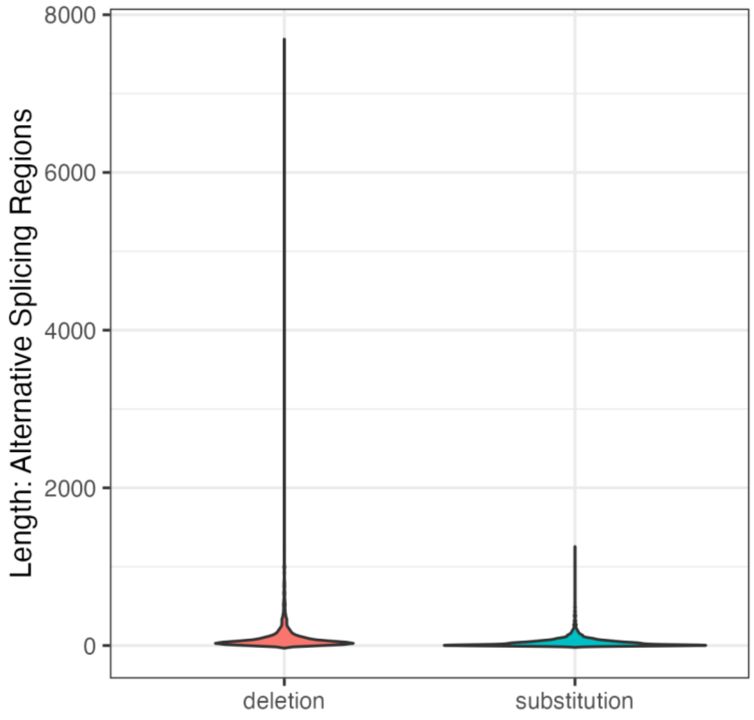


Supplemental Figure S2. Length distribution of deletion and substitution regions in canonical isoforms.

**Supplemental Tables**

Supplemental Table S1. Protein structural features.

| Feature Type | Structural Features | Feature Details |
|---|---|---|
| 8-Class Secondary Structures | $\alpha$-helix | DSSP code: H |
| | $\beta$-sheet | DSSP code: B |
| | $\beta$-strand | DSSP code: E |
| | 3-10 helix | DSSP code: G |
| | $\pi$-helix | DSSP code: I |
| | Turn | DSSP code: T |
| | Bend | DSSP code: S |
| | Loop | DSSP code: - |
| 3-Class Secondary Structures | Coils (loop, turn, bend) | DSSP code: T, S, - |
| | β-strand/sheet | DSSP code: B, E |
| | α, π, 310-helix | DSSP code: H, I, G |
| Residue Exposure Levels | Core | Relative ASA: <5% |
| | Buried | Relative ASA: 5-25% |
| | Medium-buried | Relative ASA: 25-50% |
| | Medium-exposed | Relative ASA: 50-75% |
| | Exposed | Relative ASA: > 75% |
| AF2 Residue-Level Confidence Score: pLDDT | Very high | pLDDT > 90 |
| | High | pLDDT > 70 |
| | Low | 70 > pLDDT > 50 |
| | Very low | pLDDT < 50 |
| Structural Similarity | RMSD | Root-mean-square deviation |

Supplemental Table S2. Homologs and isoforms of Septin-9 protein

| Organism | Organism Type | UniProt ID | Length | Number of Alternative Isoforms |
|---|---|---|---|---|
| Saccharomyces Cerevisiae | Fungi | P25342 | 322AA | 0 |
| Kluyveromyces Lactis | Fungi | Q6CK80 | 337AA | 0 |
| Ashbya Gossypii | Fungi | Q75ES8 | 328AA | 0 |
| Schizosaccharomyces Pombe | Fungi | Q09116 | 331AA | 0 |
| Magnaporthe Oryzae | Fungi | G4ML89 | 337AA | 0 |
| Neurospora Crassa | Fungi | Q1K4P2 | 337AA | 0 |
| Danio Rerio | Fishes | Q6TGX3 | 584AA | 0 |
| Xenopus Tropicalis | Amphibians | A0A6I8QNF1 | 576AA | 5 |
| Gallus Gallus | Birds | Q5F3T2 | 584AA | 0 |
| Mus Musculus | Mammals | Q80UG5 | 583AA | 5 |
| Rattus Norvegicus | Mammals | Q9QZR6 | 564AA | 7 |
| Homo Sapiens | Mammals | Q9UHD8 | 586AA | 26 |
| Bos Taurus | Mammals | A0A3Q1MH57 | 585AA | 5 |

Supplemental Table S3. Percentage of secondary structures in Septin-9 protein

| Organism | Organism Type | $\alpha$-helix | $\pi$-helix | 310-helix | $\beta$-strand | $\beta$-sheet | Bend | Turn | Loop |
|---|---|---|---|---|---|---|---|---|---|
| Saccharomyces cerevisiae | Fungi | 0.38 | 0.04 | 0.03 | 0.17 | 0.01 | 0.09 | 0.08 | 0.21 |
| Kluyveromyces lactis | Fungi | 0.36 | 0.04 | 0.02 | 0.16 | 0.01 | 0.07 | 0.09 | 0.25 |
| Ashbya gossypii | Fungi | 0.35 | 0.03 | 0.03 | 0.16 | 0.01 | 0.1 | 0.09 | 0.23 |
| Schizosaccharomyces pombe | Fungi | 0.33 | 0.03 | 0.02 | 0.17 | 0 | 0.09 | 0.1 | 0.26 |
| Magnaporthe oryzae | Fungi | 0.36 | 0.03 | 0.01 | 0.17 | 0 | 0.09 | 0.1 | 0.24 |
| Neurospora crassa | Fungi | 0.35 | 0.03 | 0.02 | 0.16 | 0 | 0.1 | 0.09 | 0.24 |
| Danio rerio | Fishes | 0.19 | 0.02 | 0.01 | 0.09 | 0 | 0.04 | 0.05 | 0.6 |
| Xenopus tropicalis | Amphibians | 0.2 | 0.02 | 0.01 | 0.1 | 0 | 0.04 | 0.05 | 0.59 |
| Gallus gallus | Birds | 0.23 | 0.02 | 0.02 | 0.1 | 0 | 0.05 | 0.05 | 0.53 |
| Mus musculus | Mammals | 0.19 | 0.02 | 0.01 | 0.1 | 0 | 0.03 | 0.06 | 0.59 |
| Rattus norvegicus | Mammals | 0.21 | 0.02 | 0.01 | 0.1 | 0 | 0.04 | 0.07 | 0.55 |
| Homo sapiens | Mammals | 0.19 | 0.02 | 0.01 | 0.1 | 0 | 0.04 | 0.05 | 0.59 |
| Bos taurus | Mammals | 0.19 | 0.03 | 0.01 | 0.1 | 0 | 0.03 | 0.04 | 0.6 |

Supplemental Table S4. Secondary structure changes before and after missense mutations.

| mutation | R_region | secondary_structure_wt | secondary_structure_mt |
|---|---|---|---|
| D252V | R1 | Loop | Loop |
| K257T | R1 | Loop | Loop |
| I260V | R1 | Loop | Bend |
| L266V | R1 | Turn | Turn |
| G272V | R1 | Loop | Loop |
| G273R | R1 | Loop | Turn |
| N279K | R2 | Loop | Loop |
| L284R | R2 | Loop | 3-10 Helix |
| S285R | R2 | Turn | 3-10 Helix |
| V287I | R2 | Loop | Loop |
| C291R | R2 | Loop | Loop |
| N296D | R2 | Loop | Turn |
| N296H | R2 | Loop | Turn |
| V300I | R2 | Loop | Loop |
| P301T | R2 | Loop | Loop |
| P301S | R2 | Loop | Loop |
| P301L | R2 | Loop | Loop |
| G303V | R2 | Loop | Loop |
| S305I | R3 | Loop | Loop |
| S305N | R3 | Loop | Loop |
| L315R | R3 | Loop | Loop |
| K317M | R3 | Turn | Turn |
| K317N | R3 | Turn | Turn |
| S320F | R3 | Loop | Loop |
| S320Y | R3 | Loop | Loop |
| P332S | R3 | Loop | Loop |
| G335S | R3 | Loop | Loop |
| G335A | R3 | Loop | Loop |
| G335V | R3 | Loop | Loop |
| Q336R | R4 | Loop | Loop |
| Q336H | R4 | Loop | Loop |
| V337M | R4 | Loop | Loop |
| E342V | R4 | Loop | Loop |
| S352L | R4 | Loop | Loop |
| S356T | R4 | Turn | Turn |
| I360V | R4 | Turn | Turn |
| V363I | R4 | Loop | Loop |
| P364S | R4 | Loop | Loop |
| G366R | R4 | Loop | Loop |
